# Biochemical reconstitution of a major age-related cancer mutational signature by heat-induced spontaneous deamination of 5-methylcytosine residues, repair of uracil residues, and DNA replication

**DOI:** 10.1101/2024.05.22.595323

**Authors:** Tomohiko Sugiyama

## Abstract

Non-enzymatic spontaneous deamination of 5-methylcytosine, producing thymine, is the proposed etiology of cancer mutational signature 1, which is the most predominant signature in all cancers. Here, the proposed mutational process was reconstituted using synthetic DNA and purified proteins. First, single-stranded DNA containing 5-methylcytosine at CpG context was incubated at an elevated temperature to accelerate spontaneous DNA damage. Then, the DNA was treated with uracil DNA glycosylase to remove uracil residues that were formed by deamination of cytosine. The resulting DNA was then used as a template for DNA synthesis by yeast DNA polymerase δ. The DNA products were analyzed by next-generation DNA sequencing, and mutation frequencies were quantified. The observed mutations after this process were exclusively C>T mutations at CpG context, which was very similar to signature 1. When 5-methylcytosine modification and uracil DNA glycosylase were both omitted, C>T mutations were produced on C residues in all sequence contexts, but these mutations were diminished by uracil DNA glycosylase-treatment. These results indicate that the CpG>TpG mutations were produced by the deamination of 5-methylcytosine. Additional mutations, mainly C>G, were introduced by yeast DNA polymerase ζ on the heat-damaged DNA, indicating that G residues of the templates were also damaged. However, the damage on G residues was not converted to mutations with DNA polymerase δ or ε. These results provide biochemical evidence to support that the majority of mutations in cancers are produced by ordinary DNA replication on spontaneously damaged DNA.

## Introduction

Hydrolytic deamination of cytosine (C), producing uracil (U), is one of the most common non-enzymatic decompositions of DNA bases [1]. Since U residues in cellular DNA can pair with incoming dAMP during DNA replication, unrepaired C deamination can readily cause C>T mutations. However, the vast majority of U residues in DNA are removed by a strong activity of cellular uracil DNA glycosylase (Udg) that is a part of the base-excision repair mechanism. In the human genome, about 80% of C residues in CpG context have 5-methylcytosine (5-meC) modification [2, 3]. While this modification is crucial for genomic imprinting and some specific gene regulations [4], deamination of 5-meC produces a T residue that potentially produces T-G mispairs in double stranded DNA. While T-G mispairs can be recognized by thymine DNA glycosylase (TDG) or methyl-CpG binding domain protein 4 (MBD4) [5, 6], these processes are considered less efficient than repairing U residues in DNA. Furthermore, deamination occurs more efficiently *in vitro* at 5-meC than at unmodified C [7]. Consistently, C>T at CpG sequence is the most observed spontaneous mutation in the HPRT gene of cultured human cells [8].

Massive analyses of human cancer genomes revealed multiple recurring patterns of mutations in cancers, termed cancer mutational signatures [9, 10]. Among about 60 distinct cancer signatures, the most prevalent one in all cancers (signature 1) has C>T mutation in CpG context (<u>C</u>pG > <u>T</u>). From the pattern of mutations, this signature was proposed to be caused by deamination of 5-meC [9]. Signature 1 is observed in any cancer type as well as in non-cancer tissues, and the mutations increase with the age of the sample donor [11-13]. These characteristics are consistent with the nature of spontaneous deamination of 5-meC.

Direct analyses of DNA damages in cell-free systems revealed that the spontaneous deamination of C and 5-meC can be accelerated with temperature, following first order kinetics [14-16]. The reaction occurs much faster in single-stranded DNA (ssDNA) than in double-stranded DNA (dsDNA) [17, 18]. The temperature and reaction rate have a linear relationship in the Arrhenius plot, and extrapolation of the plot indicated that an estimated half-life of C residue was 20-100 years in ssDNA at 37°C at neutral pH [7, 19, 20]. Experiments at high temperature have also identified other DNA decompositions, including depurination/depyrimidination producing abasic sites, and deamination of G and A residues producing xanthine and hypoxanthine, respectively [19]. Rates of deamination of G and A residues are 50 to 100-fold lower than that of C-deamination [19], and the contribution of these damages in spontaneous mutations has not been clear.

In this work, a process of heat-induced mutagenesis was reconstituted in a cell-free system. Synthetic ssDNA containing 5-meC was incubated at an elevated temperature, treated with Udg, and then used as a template for primer extension by yeast DNA polymerase δ (yPol δ). Direct sequencing of the DNA products identified mutations with a similar spectrum to cancer signature 1, confirming its proposed biochemical etiology. In addition, C>G mutations on damaged G residues, which did not exist in signature 1, were detected in the presence of DNA polymerase ζ. The nature of the G damage has also been investigated.

## Materials and Methods

### DNA and Proteins

Sequences of all synthetic DNA molecules used in this study have been previously published [21]. The structures of single-stranded DNA (ssDNA) templates (template A to G) and primers are shown in S1 Fig. Double-stranded DNA (dsDNA) templates were prepared by annealing ssDNA templates with the “top strands” [21] that were complementary to the “variable regions” (see Fig 1B and S1 Fig). To produce dsDNA with 5-meC modification at CpG sites, 2 μM of dsDNA was incubated with CpG methyltransferase (M. SssI, New England Biolabs) in the presence of 200 μM of S-adenosylmethionine, following the manufacturer’s protocol. To produce an ssDNA template containing 5-meC, 1 μM of methylated dsDNA and 10 μM of competitor DNA, which was complementary to the top strand, were incubated at 94°C for 10 sec and cooled to 37°C within 30 min. Then, the DNA was desalted by passing a G-25 spin column that was preequilibrated with H_2_O.

**Fig 1.**
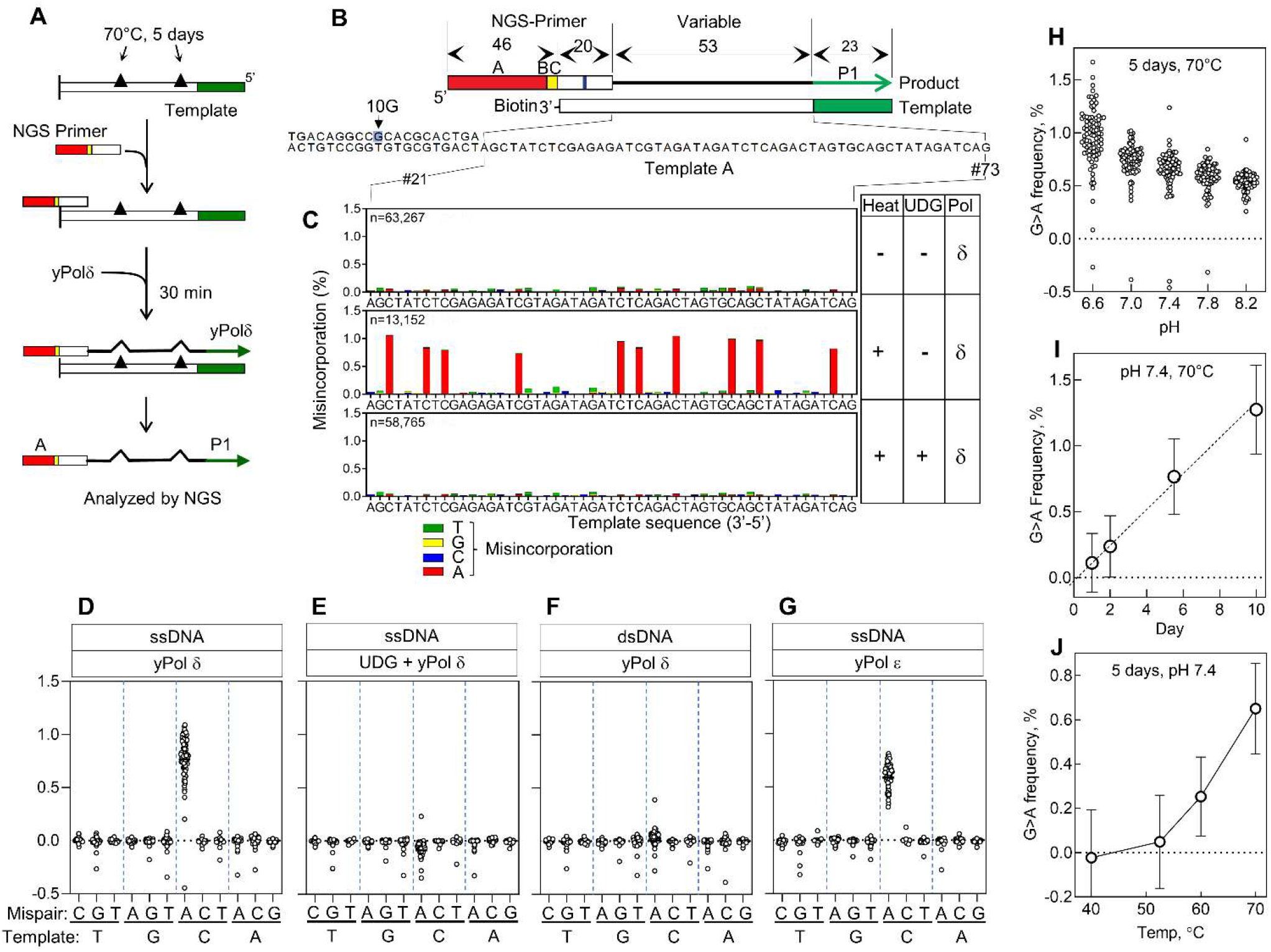
Heat-induced mutagenesis. (A, B) Illustration of the mutation assay system. The NGS adaptors (red and green bars) are located separately on the primer and template, so that only a fully extended primer can be recognized by the NGS system. To avoid extension from the template, 3’-OH of the template was blocked by biotin. To further eliminate the template-extension product from the analysis, the primer/template hybridization region contains one mismatch (10G) to select primer-extension products during the data analysis. Unique barcodes (“BC”) on the template were used to distinguish the products of separate reactions. (C) Examples of the results. Damaged or undamaged single-stranded DNA (template A) was hybridized with the primer and extended by yPol δ, and the mutation frequencies were mapped on the template sequence. In the bottom panel, the damaged DNA was treated with EcUdg before the primer extension. (D-G) Single-base substitution frequencies in total of 350-nt regions were obtained under the conditions that are indicated above each graph. Damage-induced mutation frequencies were calculated by subtracting background frequencies on undamaged templates. Bars are means. The ‘n’ values of template bases are A = 88, C = 90, G = 88, and T = 84. (H-J) The ssDNA templates were treated by heat under the standard conditions except for varying pH (H), incubation period (I), or temperature (J), and then subjected to the primer extension by yPol δ. The frequencies of G>A mutations in the products (total of 350-nt regions) are shown (I and J show mean+/-SD, n=90).

Yeast Pol δ (yPol δ: complex of Pol3-Pol31-Pol32 subunits) in which Pol32 was tagged with C-terminal His_6_ [22], yeast Pol ε (yPol ε: catalytic subunit) with C-terminal His_6_-tag [23], yeast Pol ζ (yPol ζ: complex of Rev3-Rev7-Pol31-Pol32 subunits) in which Rev7 and Pol32 were tagged with FLAG and His_6_, respectively [23], human Pol η (hPol η) with C-terminal His_6_-tag [21], human Pol ι (hPol ι) with C-terminal His_6_-tag [21], human Pol κ (hPol κ) with C-terminal His_6_-tag [21], and yeast Rev1 (yRev1) with C-terminal His_6_-tag [21], were purified as described in our previous papers. Concentrations of the polymerases were adjusted to 200 nM with 30 mM Tris-HCl(pH 7.5), 50 mM NaCl, 1 mM EDTA, 1 mM DTT and 5% glycerol, and stored at -80°C. *E. coli* Uracil DNA glycosylase (EcUdg), human AP endonuclease (hApe1), and human Smug1 (hSmug1) were purchased from New England Biolabs.

### Heat-treatment of DNA

An equimolar mixture of seven DNA templates (template A-G) were incubated in a microcentrifuge tube at 70°C for 5 days on a Pekin Elmer Thermal Cycler 460. The standard damaging reaction (40-100 μl) contained 100 nM DNA (total concentration of seven templates), 100 mM KCl, 10 mM MgCl_2_, 1 mM EDTA, and 50 mM K-hepes (pH7.4) [18], and the solution was overlayed with 100 μl of mineral oil. After the heat-damaging reaction, mineral oil was removed, and the DNA was stored at -20°C.

### Glycosylase treatment

Where indicated, DNA was treated by EcUdg (0.5 units/μl), hApe1 (1 units/μl), or hSmug1 (0.5 or 1.0 units/μl) at 37°C for 30 min in the buffer that was supplied by the manufacturer. The reactions were stopped by the phenol/chloroform/isoamyl alcohol extraction, and then the DNA was precipitated with ethanol and resuspended into the desired buffer for primer extension.

### Primer Extension for mutation assay

The primer-extension reaction for mutation assay was carried out essentially as published [21]. In the standard reaction (10 μl), the DNA template (0.10 pmol) and an NGS primer (0.1 pmol) were annealed by heating to 94°C for 4 sec and cooling to 37°C within 15 min, and yPol δ (0.2 pmol in 1 μl) was added to start primer extension. At this point, the reaction buffer contained 25 mM Tris-acetate (pH7.5), 50 mM NaCl, 4 mM MgCl_2_, 100 μg/ml BSA, 5 mM DTT, and 100 μM of each of the four dNTPs. After incubating for 30 min at 37°C, the reaction was stopped by adding 1 μl of 0.5 M EDTA, diluted 2-fold by TE buffer, and deproteinized by phenol/chloroform/isoamyl alcohol extraction. The DNA was precipitated with ethanol and resuspended into 10 μl H_2_O. Samples were then pooled and analyzed by an Ion Torrent GeneStudio S5 (ThermoFisher).

When a dsDNA template was used, the top strand was sequestered by competitor DNA (1 pmol), which was added to the reaction before the annealing step. When two polymerases were used in a single reaction, the DNA was incubated first with yPol δ (0.2 pmol in 1 μl) for 10 min, and then the second polymerase (0.2 pmol in 1 μl) was added and incubated for an additional 20 min. When hPol ι was used as a second polymerase, 200 μM of MnCl_2_ was included in the reaction.

### NGS data processing

A detailed procedure for the data processing was previously published [21]. In brief, base alterations in the NGS output sequences were mapped on the reference sequences using the Lastz 1.3.3 sequence alignment tool [24]. The mutation frequencies (% in total qualified reads) at individual bases were calculated from the Lastz output, and then trinucleotide mutation spectra were calculated by Microsoft Excel [21]. For comparison, COSMIC signatures (ver 3.2) were downloaded from https://cancer.sanger.ac.uk/cosmic/signatures. The numbers of qualified reads (n) obtained by NGS analyses are shown in S1 Table. GraphPad Prism version 9 and Microsoft Excel were used to compute statistical values. Statistical analyses of individual experiments including “n” are described in the figures and figure legends.

## Results and Discussion

### Biochemical reconstitution of heat-induced mutational processes

To analyze the mutations by heat-induced DNA damages *in vitro*, synthetic ssDNA molecules (template A-G; S1 Fig) were incubated at 70°C for 5 days and used as templates for primer extension by replicative DNA polymerases yPol δ (Fig 1A). Templates and primers were designed so that only fully extended primers were recognized by an NGS system [21]. Products of separate primer extension reactions were distinguished by unique barcodes on the primers (‘BC’ in Fig 1B). Raw sequence reads were processed so that the base-substitutions were mapped on the template sequence. Typically, 10,000 to 200,000 qualified reads were obtained for each template (‘n’ in Fig 1C and S1 Table), and the mutation frequencies were calculated from the number of base substitutions and total qualified reads. Fig 1C shows some examples of the mutation frequencies on a template (template A), showing misincorporations of dAMP (red bars) at C residues of the template, causing G>A mutations. All seven templates showed similar types of mutations (S2 Fig).

The mutation frequency data of total 350-nt regions (50-nt x 7 templates, shown in S1 Fig) are summarized in Fig 1D-G. Only G>A mutations were observed clearly above background level when replicative polymerase (yPol δ or yPol ε) were used in the reaction (Fig 1D, G). The G>A mutations were diminished when the damaged DNA was treated with *E. coli* uracil DNA glycosylase (EcUdg) before the primer extension (Fig 1E). EcUdg removes uracil from DNA, producing an abasic (AP) site. Since the replicative DNA polymerases cannot bypass AP sites [23], the reaction should not produce G>A mutations if they were caused by U residues on the templates. The G>A mutations were also diminished when dsDNA, instead of ssDNA, was incubated at 70°C (Fig 1F). All of these results support that the G>A mutations were caused by U residues, which were produced by deamination of C residues. As predicted from previous analyses of C deamination [18, 19, 25], G>A mutations accumulated linearly with incubation time and accelerated at elevated temperature and lower pH (Fig 1H-J).

Next, influences of 5-meC modification on the *in vitro* mutagenesis were analyzed in the presence and the absence of EcUdg (Fig 2). To make 5-meC modifications, the template DNA was treated with CpG methyltransferase (M.SssI). Since the methyltransferase is a dsDNA-specific enzyme, the methylation reaction was carried out with the dsDNA template, and then the top strand was removed by reannealing with the competitor DNA (Fig 2A, Materials and Methods for more details). The resulting ssDNA containing 5-meC was incubated at 70°C for 5 days to facilitate the deamination. Then the heat-treated DNA was incubated with or without EcUdg, and primer-extension with yPol δ was carried out. Finally, mutations on the fully extended products were quantified. To analyze the sequence contexts, frequencies of G>A mutations produced on C residues were organized by the trinucleotide context of the template (Fig 2B-E). In the absence of 5-meC or EcUdg, about 0.5% of G>A mutations were observed, and they were not considerably influenced by the sequence context (Fig 2B). EcUdg treatment diminished the G>A mutations on the 5-meC-free template (Fig 2C). When the 5-meC modification existed, the mutation frequencies at CpG sites were moderately increased without EcUdg treatment (Fig 2D). Side-by-side comparison showed that a 1.4 to 1.9-fold (p<0.0001) stimulation by 5-meC were observed at CpG sites, while no other site was influenced significantly (Fig 2F). This confirms previous reports showing that 5-meC is more efficiently deaminated than unmodified C, although the stimulation observed in this study is less drastic than previous reports (2-5-fold; [7, 16]). As expected, EcUdg-treatment eliminated all G>A mutations except for ones at the CpG sites if the templates had 5-meC (Fig 2E). Remaining mutations are only G>A at the CpG template context. Importantly, entire 96-dimensional mutation spectrum that is constructed under this condition (Fig 2G, H) is very similar to cancer signature 1 (Fig 2I), with a cosine similarity of 0.95 (S4 Fig). This provides a biochemical confirmation of the idea that spontaneous deamination of 5-meC followed by ordinary DNA replication, can cause equivalent mutations to the signature 1.

**Fig 2.**
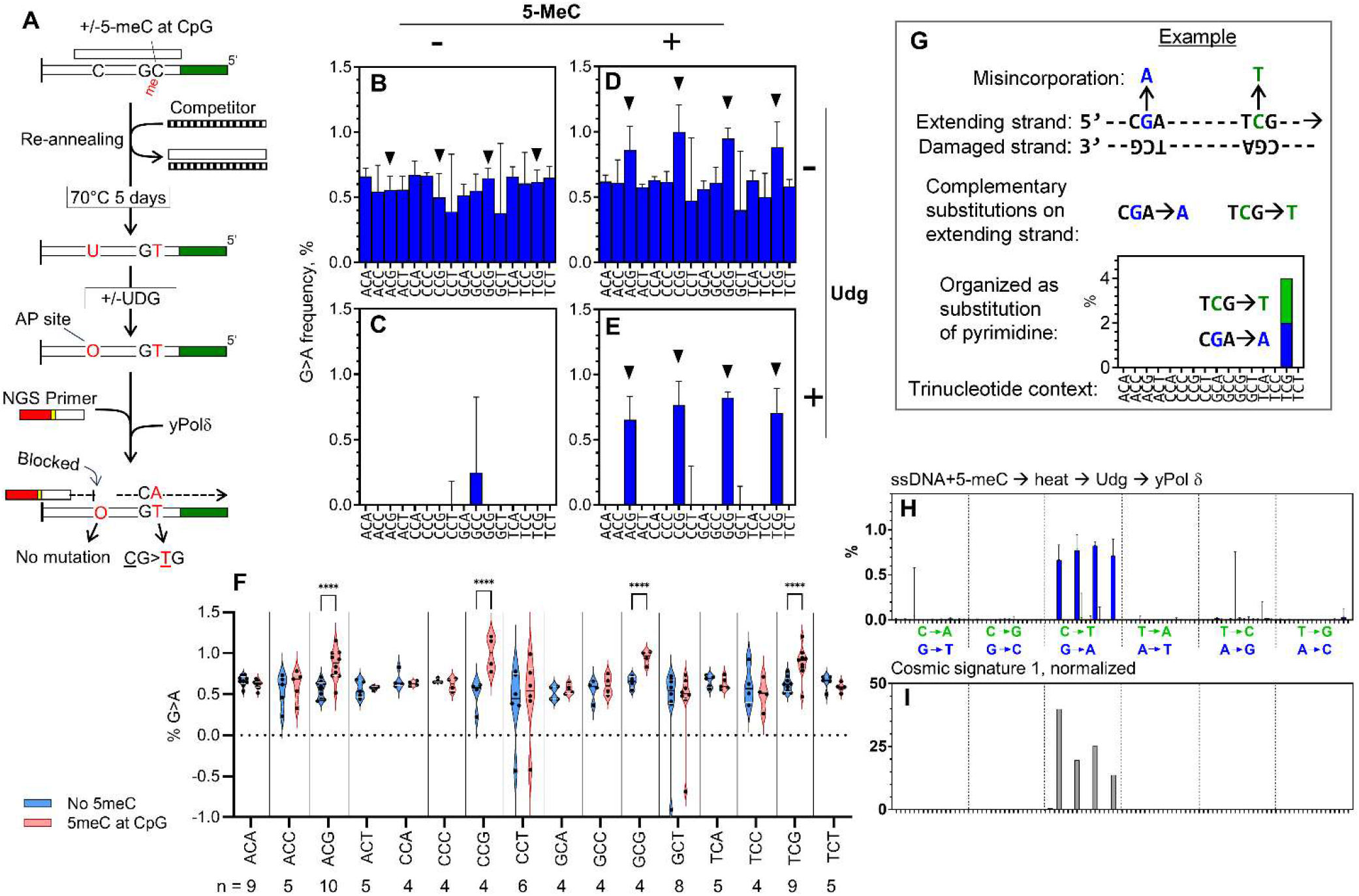
Biochemical reconstitution of cancer signature 1. (A) Illustration of heat-induced mutagenesis on 5-meC-containing DNA. (B-E) Spectra of G>A mutations on C residues of templates with different sequence contexts. The templates were modified with 5-meC and treated with EcUdg as indicated. The control DNA without 5-meC modification was prepared by the same exact procedure, except that CpG methyltransferase was omitted. Mutations on NpCpG template contexts are indicated by triangles. (F) Effects of the 5-meC modification on G>A mutation frequencies without Udg treatment were compared within each trinucleotide context. ***p<0.0001 in a 2-way ANOVA with Tukey’s multiple comparisons (n values are shown under the graph). All other pairs are scored “ns”. (G) Illustration to explain how *in vitro* mutation frequency data was organized in comparable format of cancer mutational signature. In this example, strand extension occurs from left to right on the damaged template. During the DNA synthesis, two misincorporations (G>A in blue and C>T in green) occur on the extending strand. These changes are complementary to each other including the sequence context, making them indistinguishable by genome sequencing. Since our *in vitro* approach separately quantifies such complementary mutations on the extending strand, we show them at the same position on the graph using a blue bar (purine substitution) and a green bar (pyrimidine substitution). (H) Spectrum of mutations that were produced by yPol δ on the ssDNA containing 5-meC, which was treated by heat and then by EcUdg. (I) Cancer mutational signature 1 (Alexandrov et. al, 2020) after normalization of trinucleotides appearance in human genome.

### Heat-induced guanine damage induces C>G mutation in the presence of yPol ζ

It is notable that replicative polymerase yPol δ alone can produce the signature 1-like mutation spectrum *in vitro*. It was rather unexpected to find that no other mutations were formed under these conditions, because an elevated temperature can induce multiple types of damage on DNA and some of them may be bypassed by yPol δ. It is important to note that damages not bypassed by the replicative DNA polymerases were not converted to mutations in our system. To identify the specialized polymerases that can mediate mutagenic TLS across such damages, several TLS polymerases were added to the reaction 10 min after yPol δ and incubated for an additional 20 min (Fig 3). During the first 10 min, the yPol δ-mediated primer extension was mostly completed [23]. After addition of a TLS polymerase, additional damages might be bypassed by the TLS polymerase, which may cause additional mutations (Fig 3A). Among the TLS polymerases tested (hPol η, yPol ζ, hPol κ, hPol ι, and yRev1), yPol ζ produced distinct mutations (C>G and C>A) on the G residues of the templates (Fig 3B). Especially, C>G mutations showed remarkably high frequencies (average was 0.42%), suggesting that the damaged G residue (tentatively expressed as G^#^) that caused the C>G mutation was produced at a comparable frequency to the deamination of C residues. yPol ζ-dependent C>G mutation was not observed on the heat-treated dsDNA (Fig 3B, left-bottom), indicating that G^#^ formation preferred ssDNA over dsDNA, like cytosine deamination. Other polymerases tested here did not make significant changes in the mutation spectrum, suggesting that they may not be able to bypass G^#^ or that they may bypass it without making mutations. Although this experiment could not distinguish these possibilities, some polymerase, especially hPol η, might mediate the nonmutagenic TLS in this reaction as reported [26].

**Fig 3.**
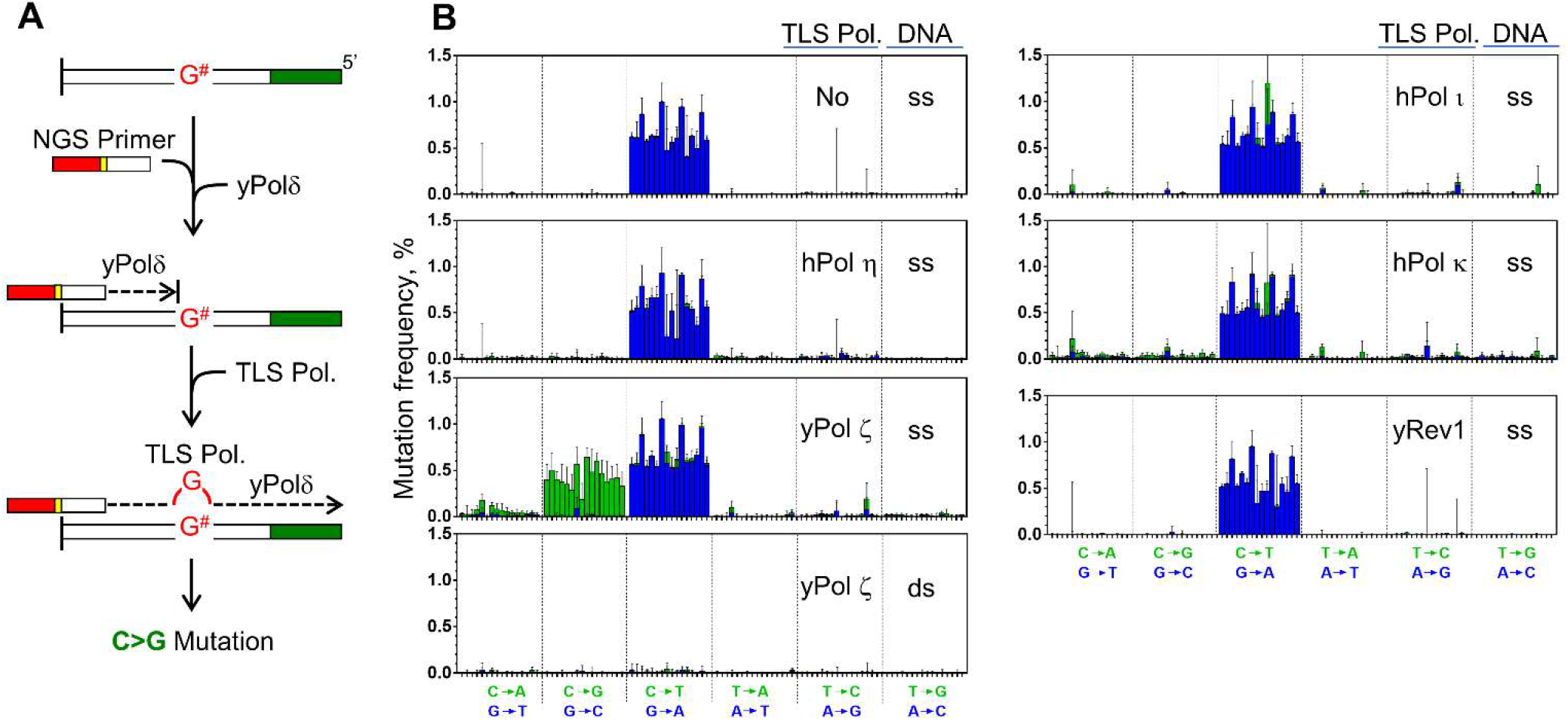
TLS-polymerase-dependent mutagenesis. (A) Illustration of the system. Heat-treated template contains unidentified damage “G^#^” that cannot be bypassed by yPol δ. TLS polymerase may be able to bypass G^#^ by inserting a mutagenic nucleotide. (B) Spectra of heat-induced mutations that were produced on the ssDNA or dsDNA by yPol δ and indicated TLS polymerases (mean +SD).

What is the G^#^ modification? Two major G modifications are known to be induced by heat at a neutral pH: deamination and base-removal, which produce xanthine and AP site, respectively [19]. To probe the identity of the G^#^, the damaged DNA was treated with various repair enzymes (Fig 4). An AP endonuclease hApe1, which cuts an AP site into a strand break, did not change C>G mutation frequencies (Fig 4C, I), indicating that the G^#^ is unlikely to be an AP site. Activity of the hApe1 can be confirmed by comparing Fig 4D and E. While yPol ζ could produce G>A mutations by bypassing AP sites that were produced by EcUdg, (Fig 4D, blue bars), all the G>A mutations were eliminated by hApe1 treatment, except for the ones on CpG sites (Fig 4E, K). This indicates that hApe1 activity was present in the reaction. The remaining major candidate of G^#^, xanthine, is cut by human Smug1 glycosylase (hSmug1), which can cut U and xanthine residues and produce strand breaks as the final products [27]. This glycosylase reduced both C>G and the G>A mutation frequencies to approximately 50% (Fig 5F, H, I), suggesting about half of the G^#^ might be xanthine. Increased amount of hSmug1 did not further reduce the mutation frequencies (Fig 4G). The remaining 50% of G^#^ could not be identified in this study. As described above, cancer signature 1 can be biochemically reproducible by the mechanism that involves spontaneous deamination of 5-meC and ordinary DNA replication. Spontaneous damages other than 5-meC deamination should occur on cellular DNA, but repairability and replicability would select damages that are eventually converted to somatic mutations.

**Fig 4.**
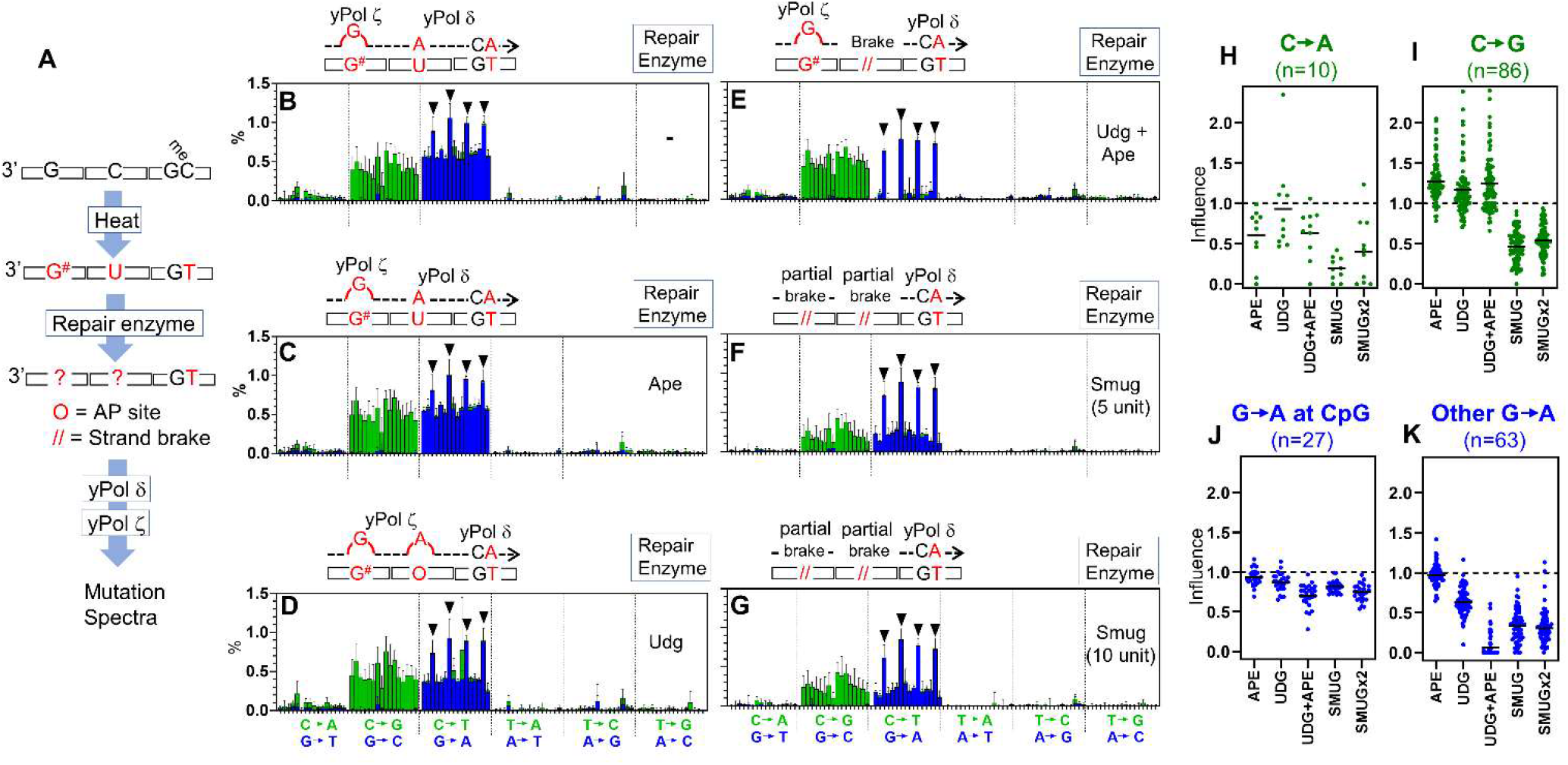
Mutation spectra in the presence of yPol ζ. (A) Illustration of the system. Heat-damaged ssDNA templates containing 5-meC were incubated with various DNA repair enzymes and then used as templates of the primer extension by yPol δ and yPol ζ as in Fig 3. (B-G) Mutation spectra made by yPol δ and yPol ζ after treatment with indicated repair enzyme. Proposed mutational mechanisms are illustrated above each graph. (H-K) Effects of glycosylases on indicated mutations were obtained as the ratios of mutation frequencies in the presence/absence of the glycosylase at individual template G and C bases. To avoid excessive data fluctuations, the ratios were calculated only at the bases that showed 0.1% or higher mutation frequencies in the absence of the glycosylase (bar = mean, n values are shown in the Figure).

## Acknowledgements

I thank Mahima Sanyal, Mya Crestwel, Elaina Grube, Eden Kenner, Noriko Kantake (Ohio University) and Sakura Sugiyama (Ohio State University) for comments on the manuscript.

## Supporting Information

**S1 Fig. Sequences of synthetic ssDNA templates (template A-G) and a NGS primer**.

**S2 Fig. The same analyses as in Fig. 1C were applied to the templates that were shown in S1**

**Fig. S3 Fig. Cosine similarity between the *in vitro* mutation spectrum (shown in Fig 2H) and normalized cancer mutational signatures**.

**S1 Table. NGS qualified read numbers**.

