## Supplemental Figures and Tables for "Biochemical reconstitution of a major age-related cancer mutational signature by heat-induced spontaneous deamination of 5-methylcytosine residues, repair of uracil residues, and DNA replication"

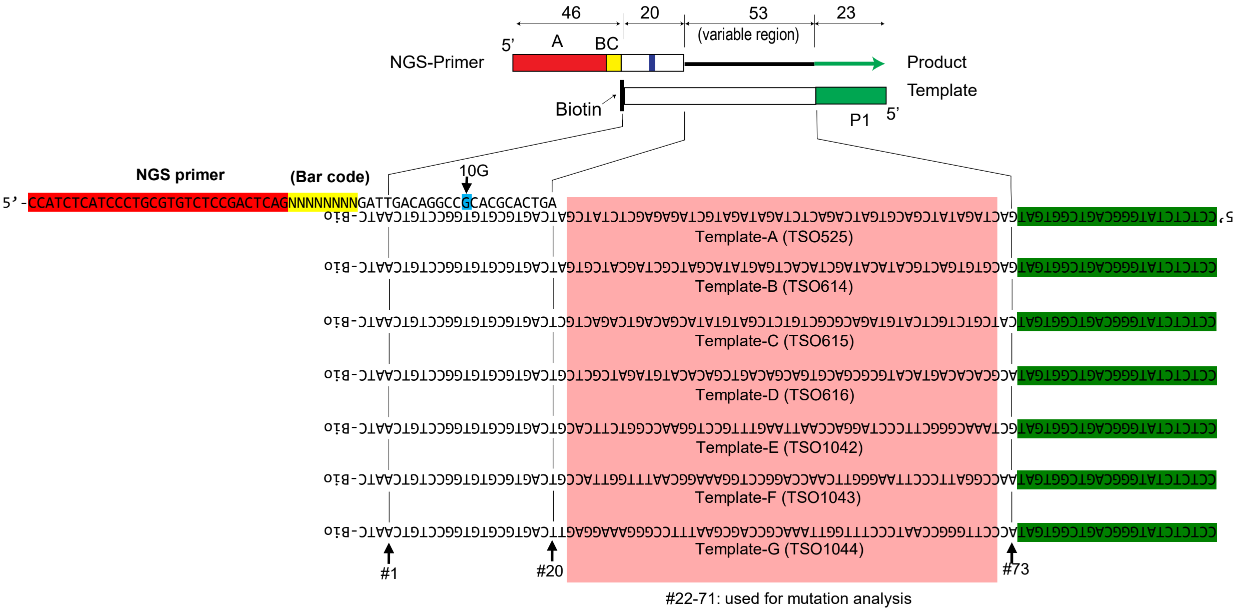


**S1 Fig Sequences of synthetic ssDNA templates (template A-G) and a NGS primer.**
All sequences were published previously (Sugiyama, et al., 2019).

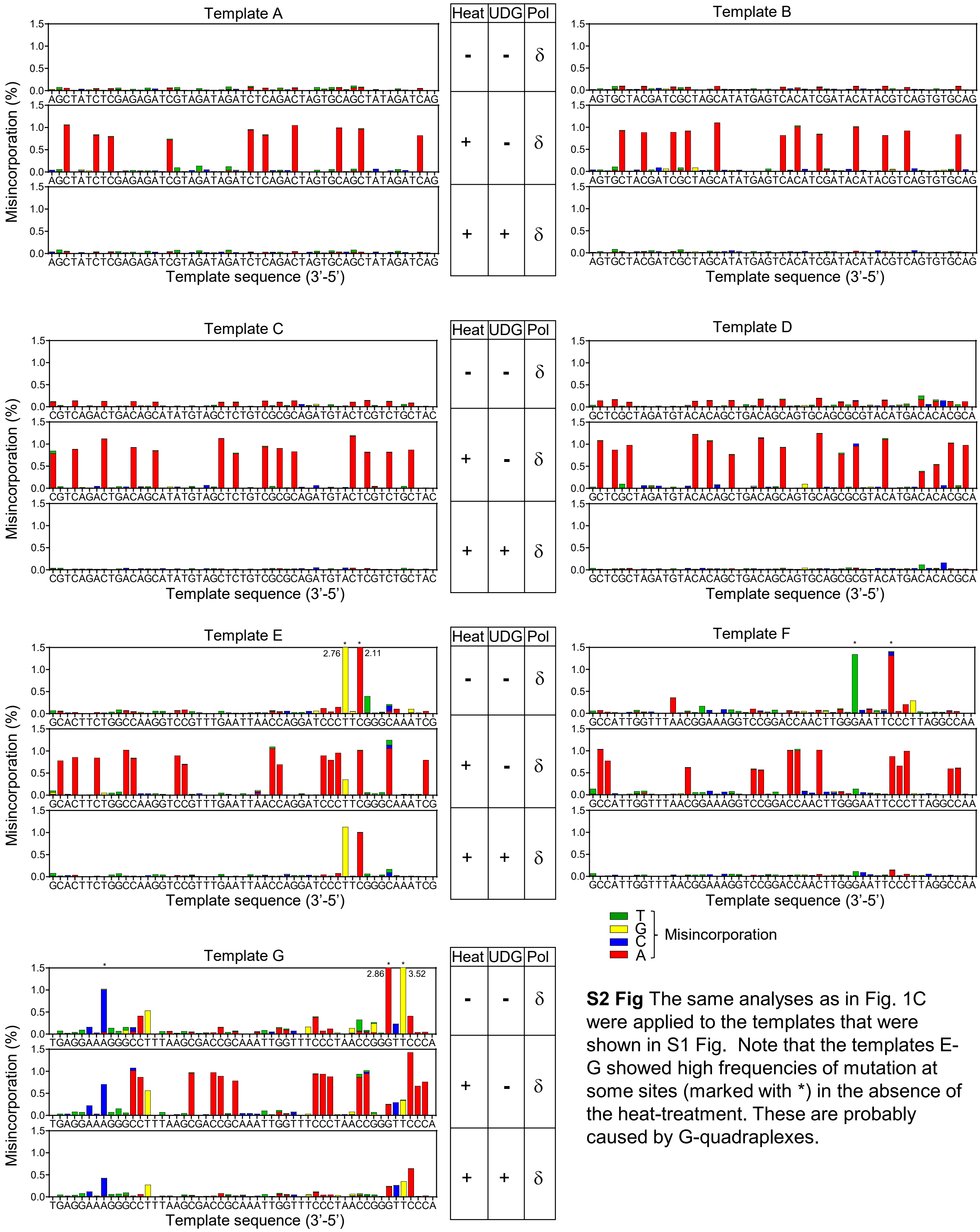


**S2 Fig** The same analyses as in Fig. 1C were applied to the templates that were shown in S1 Fig. Note that the templates E-G showed high frequencies of mutation at some sites (marked with *) in the absence of the heat-treatment. These are probably caused by G-quadraplexes.

| SIG | 1-Cos |
|---|---|
| SBS1N | 0.948956112 |
| SBS2N | 0.271035761 |
| SBS3N | 0.182293771 |
| SBS4N | 0.059045929 |
| SBS5N | 0.754802281 |
| SBS6N | 0.894633146 |
| SBS7aN | 0.405468718 |
| SBS7bN | 0.472845221 |
| SBS7cN | 0.188633991 |
| SBS7dN | 0.056080815 |
| SBS8N | 0.137234039 |
| SBS9N | 0.438840494 |
| SBS10aN | 0.142014911 |
| SBS10bN | 0.489609732 |
| SBS10cN | 0.596034793 |
| SBS10dN | 0.090068218 |
| SBS11N | 0.093896106 |
| SBS12N | 0.016417927 |
| SBS13N | 0.030242213 |
| SBS14N | 0.419166762 |
| SBS15N | 0.599932471 |
| SBS16N | 0.128680571 |
| SBS17aN | 0.030085769 |
| SBS17bN | 0.011465956 |
| SBS18N | 0.583337318 |
| SBS19N | 0.223686007 |
| SBS20N | 0.475831543 |
| SBS21N | 0.12861131 |
| SBS22N | 0.025844365 |
| SBS23N | 0.285738032 |
| SBS24N | 0.185657435 |
| SBS25N | 0.811267254 |
| SBS26N | 0.255886035 |
| SBS27N | 0.190125103 |
| SBS28N | 0.169624013 |
| SBS29N | 0.541728277 |
| SBS30N | 0.492775645 |
| SBS31N | 0.320158701 |
| SBS32N | 0.617521548 |
| SBS33N | 0.209996022 |
| SBS34N | 0.221248496 |
| SBS35N | 0.094551549 |
| SBS36N | 0.255143694 |
| SBS37N | 0.336499054 |
| SBS38N | 0.111884181 |
| SBS39N | 0.351359217 |
| SBS40N | 0.136107501 |
| SBS41N | 0.567722285 |
| SBS42N | 0.077778875 |
| SBS43N | 0.016917281 |
| SBS44N | 0.554989446 |
| SBS45N | 0.070713279 |
| SBS46N | 0.283908456 |
| SBS47N | 0.121204702 |
| SBS48N | 0.00222965 |
| SBS49N | 0.006221689 |
| SBS50N | 0.2940885 |
| SBS51N | 0.474827409 |
| SBS52N | 0.039972522 |
| SBS53N | 0.182705509 |
| SBS54N | 0.391910362 |
| SBS55N | 0.18475978 |
| SBS56N | 0.105079214 |

**S3 Fig.** Cosine similarity between the in vitro mutation spectrum (shown in Fig 2H) and normalized cancer mutational signatures (ver 3.2).
